# Chimeric 14-3-3 proteins for unravelling interactions with intrinsically disordered partners

**DOI:** 10.1101/130245

**Authors:** Nikolai N. Sluchanko, Kristina V. Tugaeva, Alfred A. Antson

## Abstract

In eukaryotes, several proteins act as “hubs”, integrating signals from a variety of interacting partners that bind to the hub through intrinsically disordered regions. Not surprisingly, one of the major hubs, the 14-3-3 protein, that plays wide-ranging roles in cellular processes, has been linked with a number of disorders including neurodegenerative diseases and cancer. A partner protein usually binds with its phosphopeptide accommodated in an amphipathic groove (AG) of 14-3-3, a promising platform for therapeutic intervention. Protein plasticity in the groove allows to accommodate a range of phosphopeptides with different sequences. So far, in spite of mammoth effort, accurate structural information has been derived only for few 14-3-3 complexes with phosphopeptide-containing proteins or various short synthetic peptides. The progress has been prevented by intrinsic disorder of partner proteins and, in case of transient interactions, by the low affinity of phosphopeptides. We reasoned that these problems could be resolved by using chimeric 14-3-3 proteins with incorporated peptide sequences. We tested this hypothesis and found that such chimeric proteins are easy to design, express, purify and crystallize. We show that when attached to the C terminus of 14-3-3 via an optimal linker, peptides become stoichiometrically phosphorylated by protein kinase A during bacterial co-expression. We determined crystal structures for complexes of chimeric 14-3-3 protein fused with three different peptides. In most of the cases, the phosphopeptide is bound inside the AG, providing invaluable information on its interaction with the protein. This approach can reinvigorate studies of 14-3-3 protein complexes, including those with otherwise challenging low affinity phosphopeptides. Furthermore, 14-3-3-phosphopeptide chimeras can be useful for the design of novel biosensors for *in vitro* and *in vivo* imaging experiments.

## INTRODUCTION

14-3-3 eukaryotic protein family is composed of abundant, medium sized proteins (∼30 kDa subunit mass) endowed with the well-known phosphopeptide-binding ability [1]. This feature allows members of the family to work in synergy with several protein kinases which, upon activation in response to specific signals, phosphorylate 14-3-3 partners triggering specific interaction with 14-3-3 proteins. This makes 14-3-3 key nodes of protein-protein interaction networks mediating a plethora of cellular processes, including apoptosis, cell division, ion channels trafficking, signal transduction, hormonal production, cytoskeleton rearrangements, etc. [1-3]. The 14-3-3 proteins have been clearly implicated in a range of human diseases including cancer and neurodegenerative disorders and as such are perspective targets for drug discovery and therapy.

In each organism, 14-3-3 proteins are usually present as several isoforms encoded by independent genes [1]. Human 14-3-3 family comprises 7 isoforms (β, σ, ζ, γ, τ, ε, η). Forming all-helical W-shape homo- and heterodimers [4-6], 14-3-3 exemplify protein modules recognizing posttranslational modifications in their partners and, with only rare exceptions [7-10], they do not interact with non-phosphorylated partners. More specifically, 14-3-3 bind protein partners having phosphoserine/phosphothreonine residues [11] and were the first phosphoserine-binding modules discovered [12]. Peptide libraries revealed the consensus motifs *I* (RSX[pS/pT]XP, where X is any amino acid) and *II* (RXY/FX[pS/pT]XP) [13] preferentially interacting with 14-3-3, implicating that protein kinases that have overlapping target sequences (e.g., AGC and CAMK protein kinase families recognizing (R/K)XXS motifs [14]) can cooperate with 14-3-3. Besides motifs *I* and *II*, the C-terminal motif *III* (pS/pTX(X)-COOH) was also found in some partners, expanding the binding repertory of 14-3-3 proteins [15]. The number of discovered 14-3-3 partners increases with time that reveals new binding sequences which significantly deviate from the “canonical” ones (motifs I-III) [16], for example, many currently known 14-3-3 partners do not have Pro/Gly in position +2 as initially believed. Significantly deviating examples include the p53 peptide LMFKpT^387^EGPD, the histone acetylase 4 peptide LPLYTSPpS^350^LPNITLGLP, the peptidylarginine deiminase isoform VI binding motif SSFYPpS^446^AEG, which are different from the motifs I-III, yet bind to the AG of 14-3-3 in conformations determined by crystallography [17-19].

At present, there are more than 2000 potential 14-3-3 interactors identified [20], explaining the involvement and important role for 14-3-3 members in many vital cellular events. Several computational tools facilitate prediction of potential 14-3-3 binding sites [20-22] and phosphopeptide binding affinities as well as contribution of distinct amino acid positions into the binding [16]. The potent phosphopeptides binding to 14-3-3 have a positively charged Arg/Lys 3 residues upstream the central phospho-residue and a downstream Gly/Pro residue conferring either flexibility or a kink at position +2, facilitating a bent conformation necessary for tight interaction in the phosphopeptide-binding amphipathic groove (AG) of 14-3-3 [13]. Remarkably, usually the equivalent non-phosphorylated sequences fail to bind to 14-3-3, pointing at a predominantly electrostatic character of interactions that initiate the primary binding [23]. Accordingly, physiologically relevant, millimolar concentrations of inorganic phosphate and sulfate may significantly inhibit 14-3-3/phosphotarget interactions by directly competing to occupy the AG site [24].

Significant finding was that 14-3-3 proteins predominantly interact with proteins enriched in intrinsically disordered protein regions (IDPRs) [25] and that phosphorylatable 14-3-3 binding sequences are mostly flexible and disordered. This poses substantial challenges for structural investigations of 14-3-3/partner interactions at atomic resolution, which is necessary for rational designing of drugs. Indeed, crystal structures for only two 14-3-3 complexes with complete protein substrates have been determined to date: the arylalkylamine N-acetyltransferase (PDB ID 1IB1 [26]) and the small heat shock protein HSPB6 (PDB ID 5LTW [27]). Limited structural information prevents out understanding of molecular processes underlying several disorders associated with the 14-3-3 proteins and ultimately slows the development of novel therapeutic approaches. Indeed, notwithstanding a wealth of solved 14-3-3/peptide complexes and recent successes in creation of various chemical stabilizers and inhibitors of some of the 14-3-3/phosphopeptide complexes [28-32], there is still a huge gap between the enormous number and variety of identified 14-3-3 interactors and accurate information about their binding. This lack of information prevents delineating a universal “14-3-3 binding law” and understanding molecular details of selectivity of 14-3-3 interaction with hundreds of competing partners. However, such information is vital for finding small molecule modifiers of a particular 14-3-3/target complex without affecting interactions of 14-3-3 with other targets. Thus, ideally, screening for small modulating compounds has to be validated using 14-3-3 complexes with the whole diverse range of peptide sequences, including low affinity peptides mediating transient interactions. However, structure determination for 14-3-3/peptide complexes is often impossible due to low affinity of peptides and/or their limited solubility preventing formation of complexes in the required 1:1 molar ratio.

All this prompted us to develop a novel approach aimed to facilitate obtaining of structural information on 14-3-3 complexes. Here we describe a simple chimeric construction of 14-3-3 proteins fused to sequences of interacting peptides that is easy to design, express in a soluble form in *Escherichia coli*, quickly and efficiently purify, and crystallize. Selected interacting peptide sequences, attached to the C terminus of 14-3-3 with a help of the optimized linker, are phosphorylated during bacterial co-expression with protein kinase A, yielding stoichiometric phosphorylation and binding of the fused phosphopeptides in the AG of 14-3-3. Crystal structures obtained are a proof-of-principle for such an approach, but also provide interesting novel structural information. The proposed approach can give an equivalent phosphopeptide conformation in the AG, but is free from limitations of a traditional co-crystallization with synthetic peptides and is compatible with high-throughput studies adequate for the wide 14-3-3 interactome. Furthermore, the approach involving 14-3-3 chimeras can accelerate the design of novel biosensors for *in vitro* screening and *in vivo* imaging studies and for construction of more complex 14-3-3-protein chimeras in future.

## RESULTS

### Design of 14-3-3 chimeras with interacting phosphopeptides

As a first prototypical chimera, we constructed a fusion of a 14-3-3 protein with a phosphopeptide belonging to the small heat shock protein HSPB6 (HSP20). Interaction between these two proteins regulates smooth muscle relaxation and the available structure of a complex with full-length HSPB6 (PDB ID 5LTW; [27]) can serve as control. We took advantage of the proximity of the natural C terminus of 14-3-3 and its amphipathic phosphopeptide-binding groove (AG) fusing the C terminus of the 14-3-3σ core segment, residues 1-231 (14-3-3σΔC), to HSPB6 peptide containing the key Ser16, which is known to be phosphorylated *in vivo* and *in vitro* by cyclic nucleotide-dependent protein kinases A (PKA) and G (PKG) [33]. Linker length between a 14-3-3 core sequence and a phosphopeptide seems crucial because it should, on one hand, be enough to reach pSer-binding pocket and, on the other hand, not interfere with the ‘normal’ binding mode of a phosphopeptide in an extended conformation. Fortunately, this was clarified thanks to the surprising asymmetric structure of a *Cryptosporidium parvum* 14-3-3, Cp14b protein, where own C-terminal peptide of Cp14b became phosphorylated upon expression in *Escherichia coli* and appeared to be bound in one of the AGs (PDB ID 3EFZ) [34] (Fig. 1A). Despite the uncommon overall fold of this rather exotic 14-3-3 member, it helped us deduce the reasonable linker length between the highly evolutionary conserved C-terminal Trp of 14-3-3 (arbitrary position 0) and the anchored phospho-residue (arbitrary position 10). Besides the structured Thr residue at arbitrary position 1, always present in the electron density maps for even C-terminally truncated 14-3-3 variants, and natural Leu residue preceding the 14-3-3 binding motif RRApS^16^APL of HSPB6, we considered GSGS linker because of its flexibility and solubility (Fig. 1B). Similar to such a prototypical 14-3-3/HSPB6 chimera CH1, we aligned and designed chimeras of 14-3-3σΔC with peptides from recently described physiological 14-3-3 partners, Gli (chimera CH2) and StARD1 (chimera CH3) (Fig. 1B), in which case the structural information, in contrast to HSPB6, is lacking. The three chimeras CH1-3 were built using the same principles and were endowed with the N-terminal His-tag cleavable by the highly specific 3C protease to facilitate their purification (Fig. 1C). To achieve stoichiometric phosphorylation of the peptides within the chimeras, we co-expressed them in *E. coli* with the catalytically active subunit of protein kinase A (PKA), known to phosphorylate all these substrates *in vivo* [33, 35, 36]. Importantly, σ-isoform of 14-3-3 is resistant to PKA phosphorylation and consequent monomerization (Tugaeva et al. 2017, submitted), mainly because it does not contain the semi-conservative interface Ser which, being phosphorylated, is reported to destabilize 14-3-3 dimers [5, 37]. In all cases, in order to alleviate crystallization, we used mutations of ^75^EEK^77^ loop residues to alanines (the so-called Clu3 mutant) by following the surface entropy reduction (SER) approach [38], which was found successful in 14-3-3/HSPB6 co-crystallization [27].

**Fig. 1.**
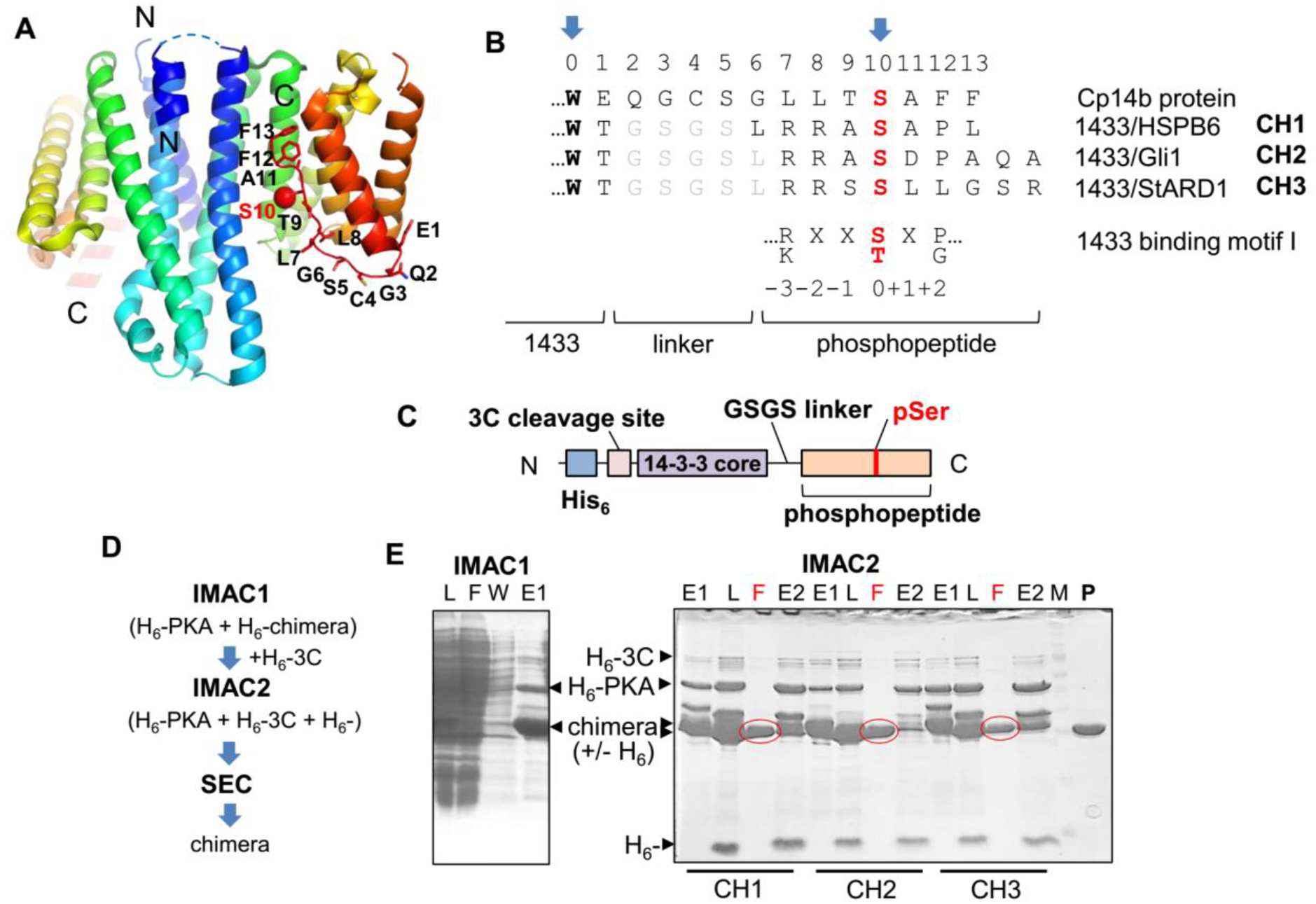
Design and production of 14-3-3 chimeras. A – Ribbon diagram of *C.parvum* 14-3-3 with phosphorylated C terminal peptide (numbered residues) bound in the AG of one of the two subunits of the 14-3-3 dimer (Cp14b protein; PDB ID 3EFZ). Each subunit is colored blue (N terminus) to red (C terminus). Drawn in PyMol 1.6.9 (Schrödinger). B – Alignment of the C-terminal region of Cp14b with designed 14-3-3 chimeras, CH1 to CH3. The conserved Trp of 14-3-3 and phosphoserine (red) are marked by arrows. Amino acid sequences linking 14-3-3 with the peptide are in grey. The 14-3-3 binding motif I is shown below the alignment. C – Schematic depiction of the proposed 14-3-3 chimeras. D – Purification scheme to obtain crystallization-quality CH proteins phosphorylated in the course of bacterial co-expression with the His-tagged PKA. Purification included subtractive immobilized metal-affinity chromatography (IMAC) for the N-terminal hexahistidine tag removal by 3C protease and size-exclusion chromatography (SEC). E, F – Electrophoretic analysis of fractions obtained during IMAC1 (only CH1 shown) and IMAC2 (CH1, CH2, CH3). L – loaded, F – flowthrough (10 mM imidazole), W – wash (10 mM imidazole), E1 – elution 1 (510 mM imidazole), E2 – elution 2 (510 mM imidazole) fractions. Note the shift of chimera bands as a result of tag removal by 3C (+/-H_6_). Flowthrough fractions (F) during IMAC2 (red circles) were polished by SEC (P – final sample) and then directly used for crystallization.

Due to the circumspect design, bacterially co-expressed and phosphorylated 14-3-3 chimeras could be easily purified by a subtractive immobilized metal-affinity chromatography (IMAC) and size-exclusion chromatography (SEC) (Fig. 1D), a procedure, in the end providing us with completely soluble (at least up to 90 mg/ml) and more than 98% pure protein samples (see Fig. 1E, lane “P”). These samples were subjected to extensive crystallization screening, while some properties of the prototypical chimera CH1 were analyzed in more detail.

### Characterization of the 14-3-3/HSPB6 protein-phosphopeptide chimera CH1

According to analytical SEC data, CH1 elution profile contained the main symmetric peak (peak “I”) corresponding to particles with an average hydrodynamic radius *R*_H_ of 3.42 nm and also a small peak (peak “II”) corresponding to particles with 4.95 nm radius (Fig. 2A). Comparison with the profiles of the monomeric mutant of 14-3-3ζ (peak at 2.77 nm, in agreement with the earlier reported value ∼2.8 nm [39, 40]) and of the 14-3-3σΔC dimer (peak at 3.56 nm) suggests that peak I of CH1 corresponds to a dimeric form, whereas peak II corresponds to a higher oligomeric form of the protein present in a much smaller amount. The smaller estimated CH1 radius (3.42 nm) relative to that of 14-3-3σΔC dimer (3.56 nm) may indicate compaction of the chimera as a result of binding of the phosphorylated heterologous C-terminal peptide within the AG. Importantly, it is believed that an unbound 14-3-3 demonstrates conformational motions along the first normal mode when its C-terminal lobes move relative to the N-terminal base of the protein, from a peptide-bound like closed to a more open conformation [6, 41]. If so, this is in agreement with the observed decrease of the average hydrodynamic radius and may be due to the phosphopeptide patching within 14-3-3 that presumably stabilizes the closed conformation, providing an easy experimental means to quickly test the correct assembly of a chimera. We can speculate that the small fraction of the larger particles with 4.95 nm appear due to the patching of a phosphopeptide from one 14-3-3 dimer to another one, giving rise to formation of tetramers (see below).

**Fig. 2.**
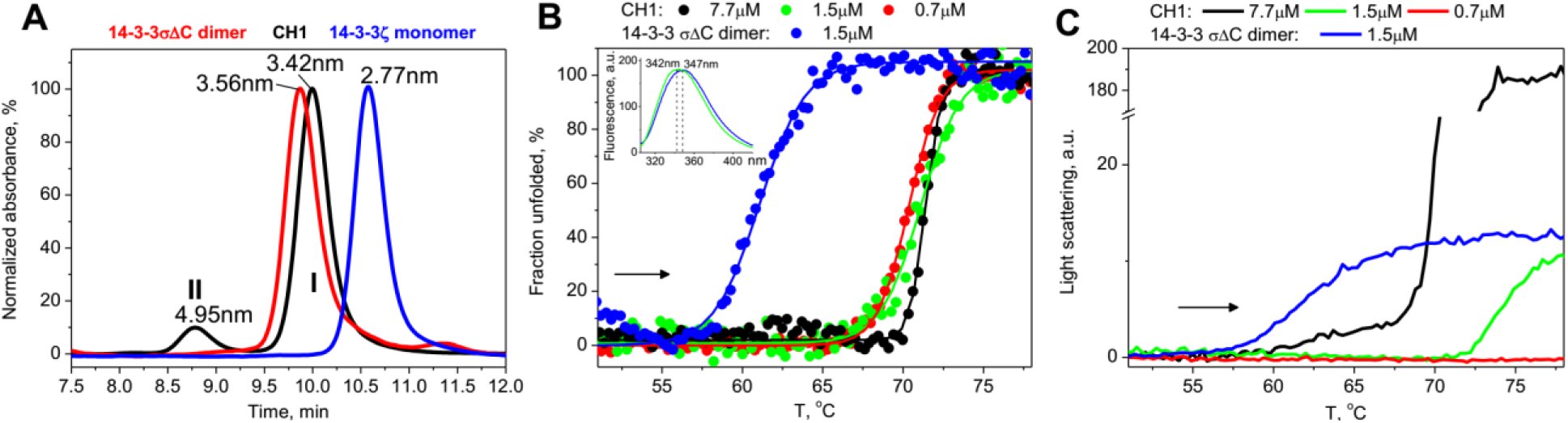
Characterization of the CH1 chimera. A – analytical SEC profiles of the monomeric mutant 14-3-3ζ [40], dimeric 14-3-3σΔC or the chimera of 14-3-3σ with the HSPB6 phosphopeptide (CH1) obtained at a 1.5 ml/min flow rate using a Superdex 200 10/300 Increase column (GE Healthcare) calibrated by proteins with known hydrodynamic radii. Elution profiles were followed at 280 nm and normalized to absorbance at maxima of the peaks. Average hydrodymanic radii corresponding to maxima of the peaks obtained from column calibration are indicated. Peaks I and II of the CH1 profile are marked. B – Intrinsic tryptophan fluorescence spectra of CH1 (green) or 14-3-3σΔC (blue) samples (1.5 μM) excited at 297 nm (slits width 5 nm) (insert) and heating of 14-3-3σΔC (1.5 μM) and CH1 (0.7-7.7 μM) samples from 10 to 80 °C at a constant rate of 1 °C/min (direction is shown by arrow) analyzed by plotting fraction unfolded against temperature [42]. C – Aggregation curves for the samples presented on panel B as temperature dependencies of light scattering accompanying aggregation.

Noteworthy, intrinsic tryptophan fluorescence measurements revealed almost 5 nm differences in the position of the maximum of CH1 spectrum (342 nm) compared to that of the 14-3-3σΔC dimer (347 nm) (Fig. 2B, insert), indicative of conformational differences in vicinity to chromophores between the two 14-3-3 variants. 14-3-3 members typically contain two native Trp residues, W59 and W230 (σ isoform numbering), with the latter situated immediately prior to the fused phosphopeptide (Fig. 1B) and therefore being potentially sensitive to structural rearrangements in vicinity. This feature may be helpful for designing of biosensors based on 14-3-3 chimeras in future (see discussion).

To further characterize CH1 chimera, we applied fluorimetry and followed changes in the intensity of intrinsic tryptophan fluorescence of CH1 or the 14-3-3σΔC dimer at two wavelengths (320 and 365 nm), transformed into temperature dependencies of the unfolded protein fraction (Fig. 2B) [42], as well as changes in light scattering during the heating of the samples at a constant rate (Fig. 2C). In this experiment, 14-3-3σ showed the main thermally-induced transition with a half-transition temperature of 61.1 °C (Fig. 2B, blue curve), followed by protein aggregation (Fig. 2C, blue curve). Under identical conditions, the half-transition temperature of CH1 was 71.1 °C, i.e. 10 degrees higher, suggesting strong stabilization of the protein, most likely as a result of phosphopeptide patching of the AG and overall compaction consistent with our SEC results. Importantly, lowering CH1 concentration down to 0.7 μM did not result in destabilization and any significant shift of the curve toward lower temperatures (half-transition temperatures only changed from 71.4 to 70.7 °C), indicative of strong phosphopeptide binding even at lowest protein concentrations (Fig. 2B). This is highly advantageous for future utilization of 14-3-3 chimeras for construction of various biosensors. Noteworthy, if added externally at equimolar protein-peptide concentration of 0.7 μM, less than 10% of the HSPB6 phosphopeptide would occupy the AG of 14-3-3 (K_D_ of the interaction is estimated as 6.3 ± 0.5 μM [27]).

Thermally-induced CH1 unfolding was accompanied by protein aggregation reflected in augmentation of light scattering as a function of temperature, however, lowering protein concentration resulted in a substantial decrease of the amplitude of the curve, and the onset of aggregation became significantly postponed (Fig. 2C). At highest protein concentration of CH1 used (7.7 μM), the aggregation curve slowly increased right after ∼55 °C, i.e. at temperature, when the unfolding and aggregation occurred in the case of the unbound 14-3-3 counterpart (Fig. 2B and C, blue lines), but in the CH1 case the sharp increase was observed only at temperatures slightly below 70 °C. This gentle rise of the scattering accompanying aggregation can be explained by gradual detachment of phosphopeptides from the AG of 14-3-3, significantly postponing the onset of the thermal transition.

### Crystal structure of the prototypical CH1 chimera

CH1 chimera crystallizes under a variety of conditions in several different crystal forms (Table 1). Thus, unlike natural C-terminal segment with intrinsically disordered parts, the peptide fusion *per se* does not hamper crystallization. One can expect that derivatives of CH1 with other phosphopeptides will crystallize equally well.

**Table 1.**
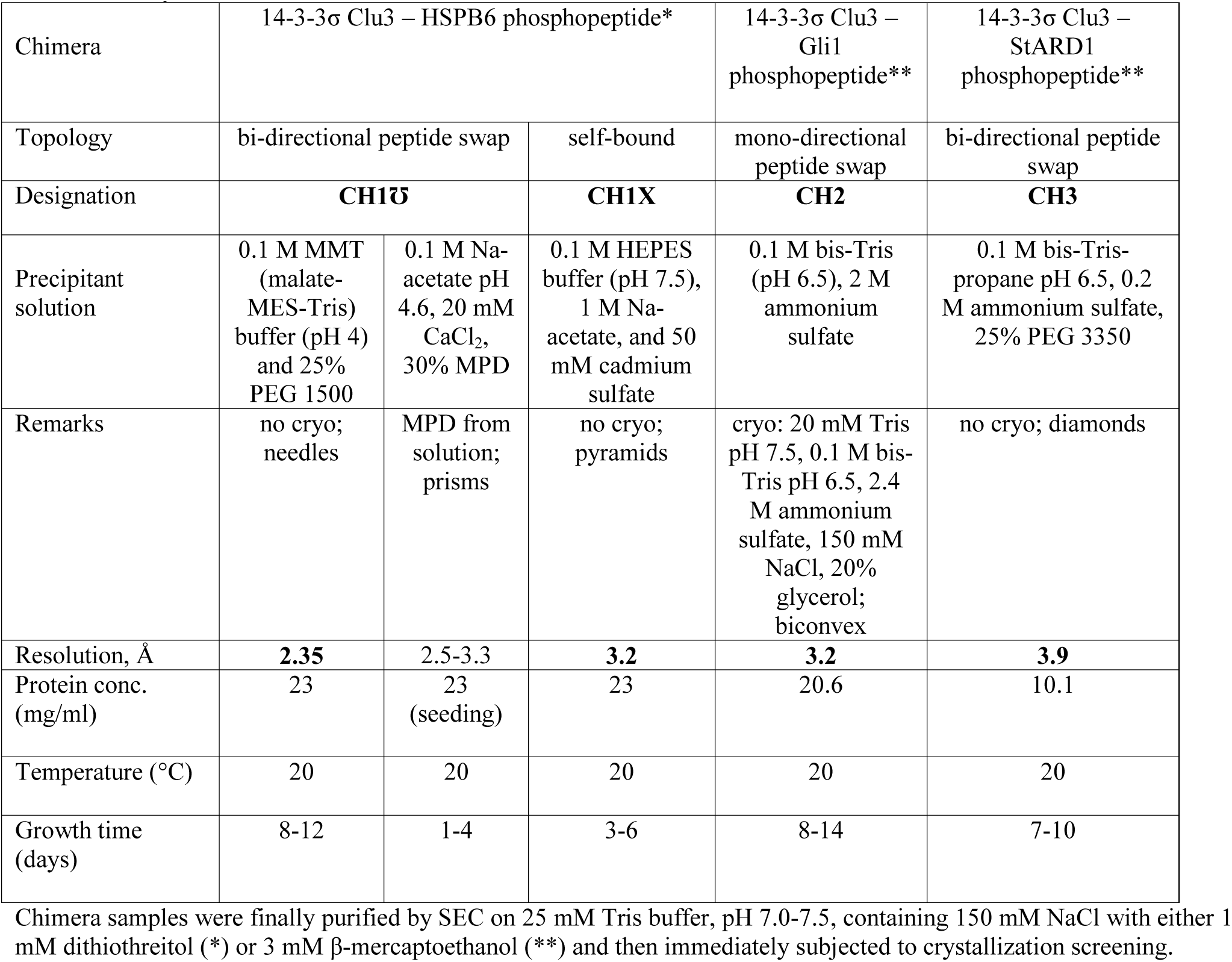
Crystallization conditions.

Two crystal forms of the CH1 chimera are remarkably distinct differing by the relative orientation and packing of 14-3-3 dimers in the crystal (Fig. 3). In one form (CH1℧, Table 1), C-terminal lobes of each of the two subunit of the 14-3-3σ dimer are in contact with the C-terminal lobes of adjacent dimers (Fig. 3A). They form an interface along the length of the 9^th^ α-helix of 14-3-3 stabilized by contacts between pairs of residues Tyr213/Tyr213 and Gln221/Gln221. As expected, the 14-3-3/HSPB6 chimeric protein produced in bacteria upon co-expression with PKA turned to be specifically phosphorylated at the authentic Ser residue in each chain (Ser16 of HSPB6 [33]). In the structure, each subunit has their phosphopeptides fused at the C terminus bound at AGs binding sites of 14-3-3 monomers from two phosphopeptides fused to the protein’s C terminus cross-patched into AGs of the adjacent monomers so that the two subunits from different 14-3-3 dimers were significantly stabilized by a subunit peptide swap. The best among three isomorphous crystals, detailed electron density map built to a 2.35 Å resolution (Fig. 3B and Table 2) allowed to easily trace the whole range of the C terminus of the CH1 chimera, including all residues of the linker, however, the very terminal Leu residue in +3 position relative to pSer16, lying just outside the primary 14-3-3 binding motif, RXXpSXP, either was absent from the map or was adopting different conformations. Surprisingly, although being very short, the GSGS linker was enough to allow phosphopeptide binding to the other’s 14-3-3 subunit.

**Fig. 3.**
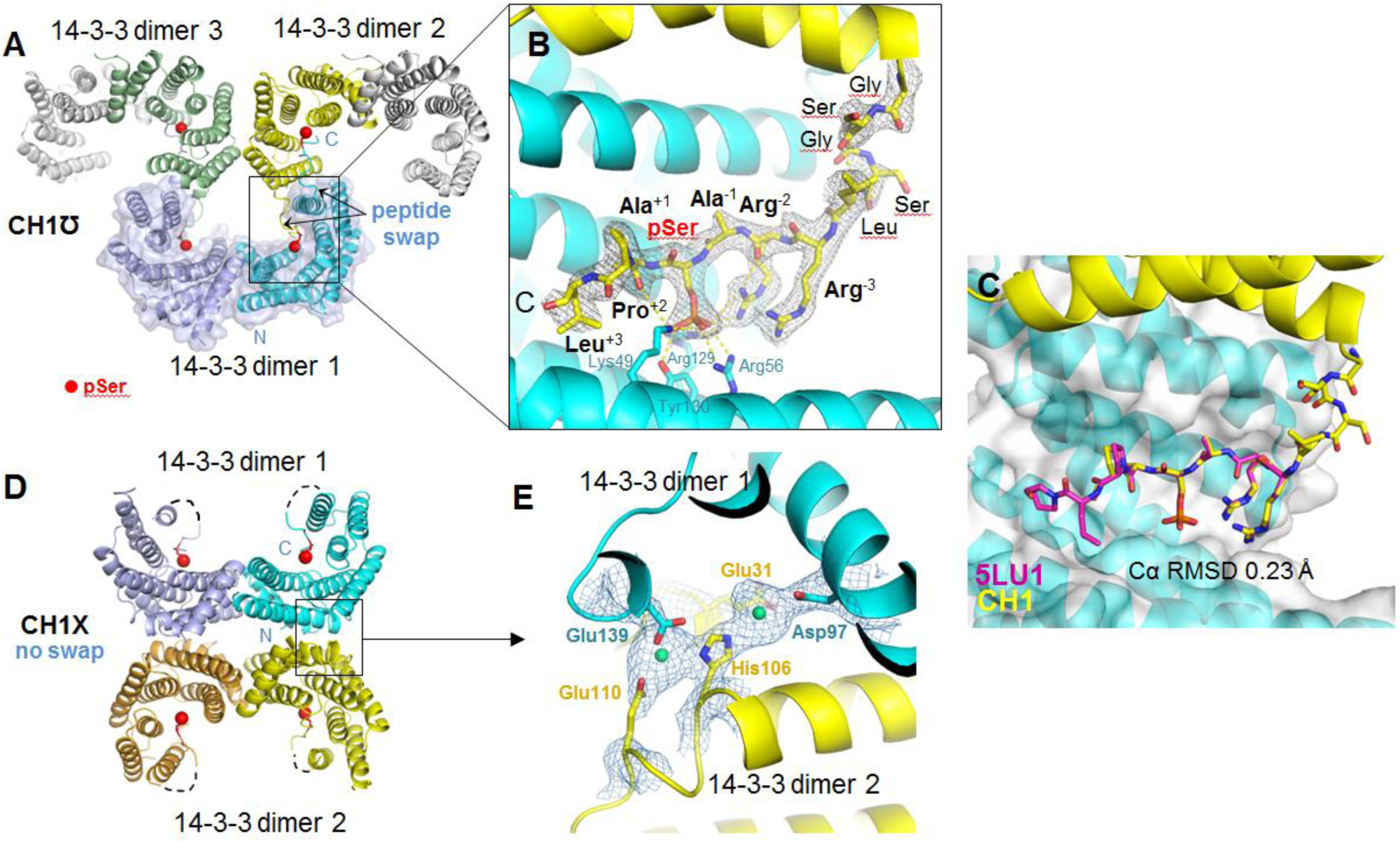
Crystal structures CH1℧ and CH1X of the prototypic chimera CH1 of 14-3-3σ with the HSPB6 peptide. A – overall organization of the ASU of the CH1℧ structure at 2.35 Å (colored 14-3-3 subunits (ribbons) forming the upturned O shape, one physiological 14-3-3 dimer is shown by semitransparent surface) with the phosphopeptide swap between unrelated 14-3-3 subunits (pSer residues are shown by red spheres). B – up-scaled view on the bound phosphopeptide conformation with the 2F_o_-F_c_ map contoured at 1σ and resolved residues of the linker or the peptide (bold font with positions relative to pSer indicated). C – Comparison of phosphopeptide conformations in the CH1 structure and 5LU1 structure with the synthetic HSPB6 phosphopeptide co-crystallized with 14-3-3σ [27]. D – overall organization of the ASU of the CH1X structure at 3.2 Å with no peptide swap (dashed lines correspond to unresolved parts of the linker). E – up-scaled view on an interface between the two 14-3-3 dimers in CH1X showing the tentative metal-binding centers (metal ions, interpreted as Cd from crystallization condition, are indicated in green) with the 2F_o_-F_c_ map contoured at 1.5σ. Molecular graphics were performed with PyMol 1.6.9 (Schrödinger).

**Table 2.** X-ray data collection and refinement statistics.

|  | CH1U | CH1X | CH2 | CH3 |
| --- | --- | --- | --- | --- |
| **Data collection** |  |  |  |  |
| Space group | $P 1 2_1 1$ | $P 2_1 2_1 2_1$ | $P 6_4 2 2$ | $P 4_1 2_1 2$ |
| Cell dimensions: $a, b, c$ (Å) | 63.6, 140.6, 68.7 | 77.4, 97.8, 158.8 | 110.4, 110.4, 174.1 | 123.3, 123.3, 162.4 |
| $\alpha, \beta, \gamma$ (°) | 90, 114.8, 90 | 90, 90, 90 | 90, 90, 120 | 90, 90, 90 |
| Resolution range (Å)* | **47 - 2.35**<br>[47-6.6] (2.51-2.35) | **46.5 - 3.2**<br>[46.5-6.9] (3.31-3.20) | **48 - 3.2**<br>[48-9.4] (3.38-3.19) | **49 - 3.9**<br>[49-11.4] (4.12-3.89) |
| Wavelength (Å) | 0.9795 | 0.9795 | 0.9795 | 0.9282 |
| $R_{\text{merge}}$** | 0.19<br>[0.07] (1.2) | 0.45<br>[0.08] (3.1) | 0.23<br>[0.03] (4.3) | 0.3<br>[0.06] (2.9) |
| $R_{\text{meas}}$ | 0.2<br>[0.08] (1.4) | 0.49<br>[0.08] (3.4) | 0.23<br>[0.03] (4.4) | 0.3<br>[0.06] (3.1) |
| $\langle I / \sigma \rangle$ | 6.5 (1.2) | 4.4 (0.7) | 14.3 (0.91) | 6.8 (1.0) |
| $CC_{1/2}$ | 0.99 (0.5) | 0.99 (0.3) | 1.00 (0.5) | 1.00 (0.3) |
| Completeness (%) | 95.5 (84.6) | 99.8 (99.9) | 99.6 (98.2) | 99.6 (98.4) |
| Redundancy | 3.9 (3.8) | 6.5 (6.7) | 23.0 (22.3) | 12.8 (12.7) |
| **Refinement** |  |  |  |  |
| No. of reflections: total | 43838 | 20548 | 10910 | 12947 |
| ‘free’ set | 1385 | 1016 | 977 | 1049 |
| $R_{\text{work}}$(%) | 19.1 | 24.7 | 21.5 | 20.9 |
| $R_{\text{free}}$(%) | 24.0 | 27.9 | 26.0 | 24.8 |
| No. of 2:2 complexes /asu | 2 | 2 | 1 | 2 |
| No. of non-H atoms: protein/ligands/solvent | 7503 / 35 / 493 | 7142 / 17 / 21 | 3607 / 20 / 9 | 7234 / 10 / 1 |
| R.m.s.d. bond lengths (Å)/ angles (°) | 0.010 / 1.0 | 0.010 / 1.0 | 0.010 / 1.1 | 0.010 / 1.1 |
| Ramachandran favoured/ outliers (%) | 97.7 / 0.1 | 98.1 / 0.1 | 96.0 / 0.4 | 96 / 0.6 |
| Molprobity score/Clash score | 1.3 / 0.99 | 1.6 / 1.05 | 1.9 / 2.07 | 2.1 / 3.04 |
| PDB code |  |  |  |  |

\*Statistics for the lowest and highest resolution shells are indicated in square brackets and parentheses, respectively
\*\*All statistics as defined in XSCALE [46].

Importantly, irrespective of the peptide swap, the phosphopeptide orientation and conformation were identical to that of the synthetic HSPB6 peptide co-crystallized with 14-3-3σ (PDB ID 5LU1 and 5LU2 [27]), with C_α_ r.m.s.d. of 0.23 Å for the residues RRApSAP (Fig. 3C) indicating highly specific binding and the absence of steric hindrances caused by chimeric construction. Consistent with many available 14-3-3/peptide crystal structures, in our CH1℧ structure the phosphate moiety forms salt bridges with the conserved 14-3-3σ residues Lys49, Arg56, and Arg129 and is H-bonded to Tyr130 (Fig. 3C). The main-chain atoms of the Ala residues immediately adjacent to the phosphoserine are H-bonded to 14-3-3 residues Asn226 and Asn175, respectively, recapitulating what was observed in 5LU1 structure [27]. This suggests that the proposed chimeras can in principle be used to get structural insights into 14-3-3 interaction with a phosphopartner without the necessity to synthesize peptides and tediously adjust protein-peptide ratios that may either interfere with crystallization or result in problems with occupancy, especially for peptides bound with moderate-to low affinity.

This important conclusion was confirmed by another crystal form of CH1, where in ASU we found two 14-3-3σ dimers located “back-to-back” (CH1X, Fig. 3D and Table 2). In this case, fused phosphopeptides apparently bound back to AG of the same subunits resulting in the fully occupied AGs. Notwithstanding lower resolution of the corresponding crystal structure (3.2 Å), all main phosphopeptide residues and some linker residues were resolved, yielding substantially the same phosphopeptide conformation as in the CH1℧ case.

Curiously, we found that the two 14-3-3 dimers in CH1X were stabilized by eight centers of the well-defined electron density located between Asp139 and Asp97 of one 14-3-3 dimer, and Glu110, His106, and Glu31 residues of the other 14-3-3 dimer in a staggered order, so that, due to the antiparallel organization of 14-3-3, four such regions could be found per protein subunit (Fig. 3E). Given that these almost identical centers of the electron density were repeated four times per CH1X and that they were coordinated by charged residues Asp/Glu and His, we assumed that there should be metal atoms (most likely, Cd which was present in crystallization media, See Table 1). Although divalent metal ions have been previously reported to bind plant members of the 14-3-3 family [43] and were even found in several 14-3-3 structures, they had different spontaneous positions, and the combination of residues Glu31, Asp97, His106, Glu110, and Asp139 was not directly linked to the ability of 14-3-3 coordinate metal ions (Glu31 and His106 coordinate Mg ion in PDB ID 3SMK). Moreover, this set seems to be unique in σ-isoform of human 14-3-3. Therefore, our structure sheds light on the remarkable tentative metal-binding property of 14-3-3 warranting further detailed exploration.

The crystal structures of CH1 also helped confirm our conclusions drawn from the SEC data. Hydrodynamics calculations in WinHydroPro program [44] using the crystallographic CH1 dimer (Fig. 3D) or dimer of dimers stabilized by reciprocal phosphopeptide patching (Fig. 3A) resulted in *R*_H_ values 34.6 and 49.8 Å, respectively, in an excellent agreement with the SEC-derived values (3.42 nm and 4.95 nm; see Fig. 2A).

### Crystal structures of CH2 and CH3 chimeras

Having found that the prototypic chimera CH1 provided equivalent structural information as if phosphopeptide was co-crystallized with 14-3-3, but with much easier workflow (protein cloning→expression/purification→crystallization), we created two other 14-3-3 chimeras, CH2 and CH3, attempting to get novel structural information by the proposed approach.

PKA-dependent phosphorylation and interaction of the transcription factor Gli (having the motif RRAS^640^DPAQA conserved in all Gli proteins, Gli1, Gli2, and Gli3), a central player in the Hedgehog signaling [35], and steroidogenic acute regulatory protein StARD1 (having motifs RRSS^57^LLGSR and RRGS^195^TCVLA) [36, 45] with 14-3-3 proteins were identified recently, however, structural details of these interactions were not clearly elucidated. For chimera construction, we chose the main phospho-site of Gli (CH2) and the first potential 14-3-3 binding motif of StARD1 (CH3) which is also found around Ser87 in human Bcl-2-like protein 11, or BimEL. CH2 and CH3 proteins were co-expressed with PKA and purified in a fully soluble form exactly as the CH1 protein, indicating that high inherent solubility of 14-3-3 is not affected by the extra C-terminal additions. Both proteins readily crystallized in various conditions giving diffraction-quality crystals of different morphology straight from commercial screens (see Table 1).

Supported by two isomorphous crystals, CH2 crystal structure solved to 3.2 Å resolution (Table 2) showed per ASU one 14-3-3 dimer mutually donating fused phosphopeptide with its symmetry mate, together forming an overall closed cyclic structure (Fig. 4A). Phosphopeptide is anchored in the AG of 14-3-3 in an expected manner and orientation, with the central Ser640 bound by the same canonical residues as in case of CH1 (Fig. 3B) and other 14-3-3/phosphopeptide complexes. Interestingly, this binding fashion leaves two opposite AGs unoccupied by two remaining phosphopeptides which are completely unresolved, however, these AGs appear to be occupied by sulfate anions present in crystallization media in high concentration (Fig. 4A and Table 1). One can speculate that very high sulfate concentration (2 M) forced phosphopeptides to partially dissociate from the AG facilitating this distinct crystal assembly, in agreement with the modulatory effect of phosphate and sulfate ions on 14-3-3/phosphotarget interactions observed *in vitro* [24]. Electron density map allowed us to readily trace all the residues of the linker and of the Gli-derived phosphopeptide, except for the last one Ala residue in position +5 from pSer^640^ (Fig. 4B). Curiously, two nearest phosphopeptide-binding 14-3-3 subunits happened to form a Cys^38^-Cys^38^ bridge accounting for a rather dense crystal packing, however, even this factor did not interfere with the conformation of the bound phosphopeptide. Thus, as a very important advantage, our approach utilizing 14-3-3 chimeras allows easy protein crystallization in different crystal forms which provide necessary structural information about the binding of a target phosphopeptide.

**Fig. 4.**
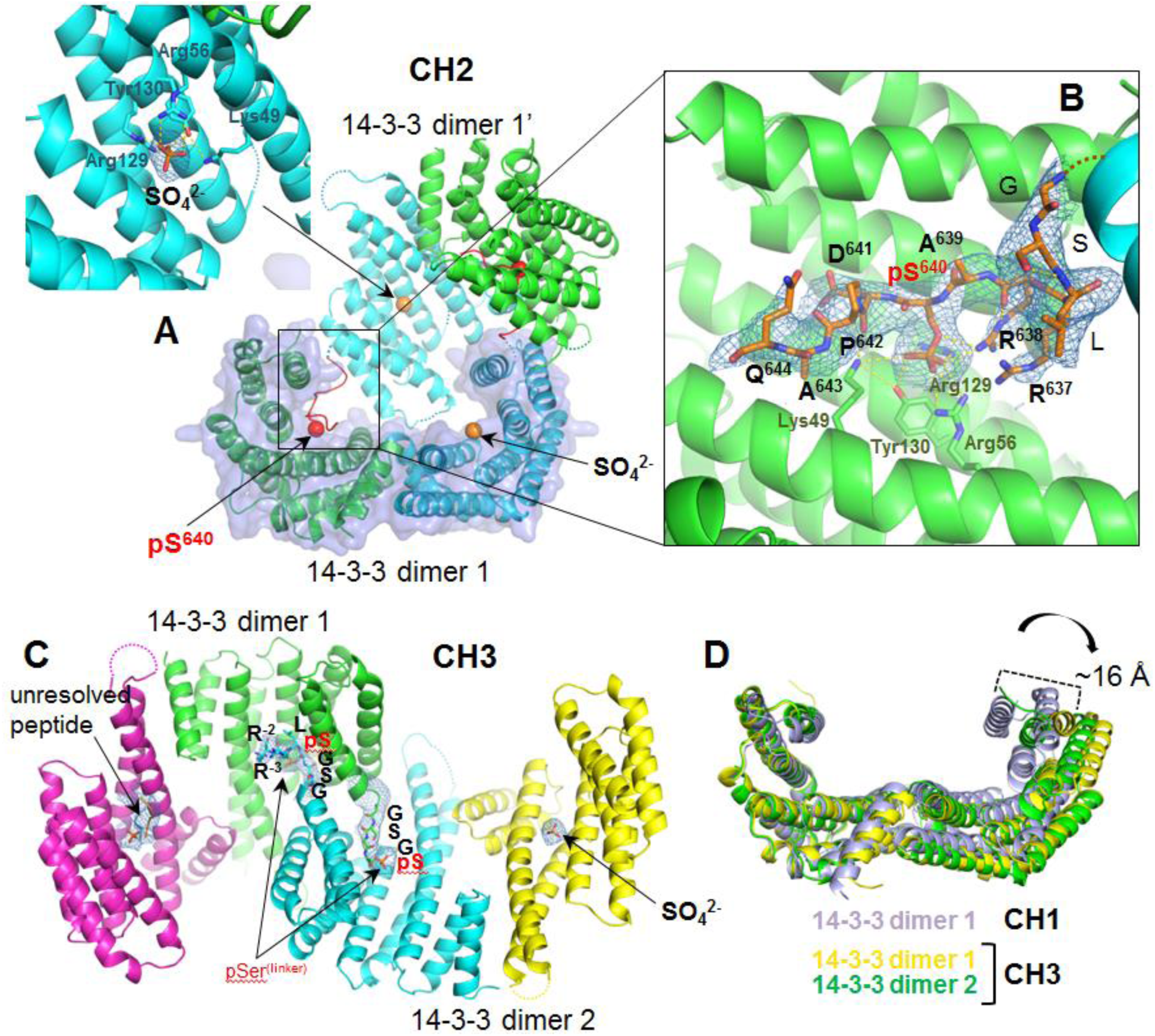
Crystal structures of the chimeras CH2 and CH3. A – overall view on 14-3-3σ chimera with Gli phosphopeptide (CH2 structure at 3.2 Å resolution) with one 14-3-3 dimer in ASU (semi-transparent surface) mutually donating its fused phosphopeptide to the symmetry mate (14-3-3 dimer 1’) and a close-up view of the sulfate anion (orange) bound in the AG of one type of the chains (inset). B – Close-up view on the bound Gli phosphopeptide with 2F_o_-F_c_ map contoured at 1σ and key residues indicated. C – overall view on 14-3-3σ chimera with StARD1/BimEL phosphopeptide (CH3 structure at 3.9 Å resolution), with two 14-3-3 dimers in ASU and regions of the 2F_o_-F_c_ map contoured at 1σ. Unresolved phosphopeptide, sulfate anion, and the phosphorylated linker GSGpSLRR are demonstrated. D – superposition of different 14-3-3 dimers by one monomer, demonstrating conformational changes from fully closed to significantly open state with ∼16-Å amplitude. Molecular graphics were performed with PyMol 1.6.9 (Schrödinger).

Simultaneously, we obtained unexpected results in the case of the CH3 chimera. Most of the crystals were poorly diffracting and we could solve only one structure to 3.9 Å resolution, with two interacting chimera dimers in ASU (Fig. 4C and Table 2). We then found that other two crystals, diffracting to a lower resolution, are isomorphous. One of the 14-3-3 chains was occupied by sulfate anion from crystallization media, and another one demonstrated density obviously larger than for phosphate/sulfate anion, likely suggesting a phosphopeptide probably belonging to the second chain of the symmetry mating 14-3-3 dimer 2 is bound, however, we failed to trace its residues due to the poor electron density in this region. Electron density map of two neighboring unrelated chimera subunits was clear enough to continue the C terminus to the counterpart’s AG. Although the distance from the C-terminal Trp residue to the positively charged residues, typically coordinating a phosphoserine, was too short to allow canonical binding, the binding of a phosphorylated residue did occur (Fig. 4C). In both cross-patching peptides, we surprisingly found that it was the second Ser residue of the linker (-GSGpS-) that became phosphorylated and bound in the AG (Fig. 4C). In one of the chains we could trace three residues beyond this pSer, LRR, indicating completely reverse orientation of the bound peptide. This clearly exemplified the extremely rare case when a peptide had been phosphorylated by PKA in a wrong orientation (N-_‥_pSXXR‥-C instead of N-_‥_RXXpS‥-C) and then was erroneously recognized by the AG of 14-3-3 in the likewise, wrong orientation. This may have been an artifact caused by relatively high levels of co-expressed PKA reducing specificity of its action, although, surprisingly, under the same conditions this did not happen to CH1 and CH2, where the same linker remained unphosphorylated. It is possible that a peptide sequence in CH3 influenced such an unspecific phosphorylation. We do not know the phosphorylation status of the target Ser and its neighboring Ser residue (RR<u>SS</u>LLGSR, underlined), however, to exclude such situations in future, Ser residues in the linker GSGS could be avoided. Although precluding us from observing the correct bound conformation of the StARD1/BimEL phosphopeptide, this unique crystal structure illustrates that the selectivity of a 14-3-3 groove is limited, at least partially explaining multiple exceptions from the known binding rules based on motifs I-III [13, 15]. This circumstance further favors the utilization of 14-3-3 chimeras to structurally study rather weak binding phosphopeptides which significantly differ from the canonical motifs that would otherwise be almost impossible because of high corresponding dissociation constants and problems with peptide atoms occupancy.

Interestingly, in CH3 structure, both 14-3-3 dimers adopted significantly more open conformation than in the CH1 or CH2 case (Fig. 4D). When superposed by one 14-3-3 monomer, the marginal states of the C-terminal α-helix of the other monomer differed by about 16 Å (dimer 1 of CH3 had an intermediate position), reminiscent of the rather unique structure of apo-14-3-3β isoform with one closed and one open subunits [6]. Surprisingly, in our case, this opening of 14-3-3σ subunit did not prevent binding of sulfate ion or a phosphorylated peptide (on Fig. 4C, the most open subunit is yellow). The observation of 14-3-3 subunits in conformations differing by the degree of subunit ‘openness’ supports such conformational rearrangements detected by molecular dynamics simulations of 14-3-3σ [41].

## DISCUSSION

The proposed approach provides a convenient means to obtain structural information on complexes of 14-3-3 with different peptides opening up new perspectives in the 14-3-3 research.

One of the advantages is that the proposed 14-3-3 chimeras are easy to design and produce in a soluble form in *E. coli* cells, when solubility is conferred by highly soluble 14-3-3 proteins and phosphorylation is achieved by a co-expressed protein kinase. PKA, used in this work for co-expression, may be substituted by another kinase which is known to phosphorylate a desired 14-3-3 binding site, provided that it is subcloned into a compatible expression vector and is soluble in *E. coli*. Gentle engineering of 14-3-3 sites to make them phosphorylatable by PKA is also an option. Alternatively, *in vitro* phosphorylation of purified 14-3-3 chimeras by commercially available protein kinase(s) may be useful for some applications.

The proposed purification procedure is affordable and straightforward comprising i) separation of a His-tagged 14-3-3 chimera with His-tagged PKA by IMAC1, ii) His-tag removal from a 14-3-3 chimera by highly specific His-tagged 3C protease (3C may be replaced by another commonly used specific protease), iii) separation of a tag-free 14-3-3 chimera from PKA and 3C during IMAC2, iv) polishing of the sample by SEC. Double IMAC steps (gradients are not necessary) allow to avoid extensive optimization of purification conditions, and the resulting samples are highly pure (>98%), soluble at tens mg/ml, monodisperse, and therefore instantly ready for crystallization. 14-3-3 chimeras readily crystallize in a variety of crystal forms, straight from commercial screens. These advantages make our approach adaptable for high-throughput studies, for example, screening of novel 14-3-3 protein interactors, validation of newly identified protein-protein interactions involving 14-3-3, and screening of small molecule modulators of the established 14-3-3/phosphotarget complexes (using already obtained crystals of a particular 14-3-3 chimera or biosensors based on 14-3-3 chimera, see below).

Substantial and obvious advantage of chimeric 14-3-3-peptide constructions is that a covalently bound peptide ensures constant 1:1 stoichiometry, which, in contrast to traditionally utilized synthetic peptides that can be fragile and of limited solubility, eliminates manipulations with the protein-peptide molar ratio for co-crystallization [27], where a large excess of a peptide may interfere with crystallization whereas a small excess may result in a partial occupancy of the AG of 14-3-3. This consideration is especially important for weak, transiently binding peptides which are apparently underexplored due to this very reason. Fusion of such peptides to 14-3-3 presents an exclusive opportunity to obtain corresponding structural information about their conformation in the AG of 14-3-3.

The optimal linker length, often an Achilles’ heel in fusion proteins, was determined in this study on the basis of the exotic crystal structure of the parasitic 14-3-3 protein Cp14b with own bound phosphorylated C terminus (Fig. 1A), and led to obtaining several crystal structures in our study (Fig. 3 and 4). It should be mentioned that despite 14-3-3ζ chimera with a pseudophosphorylated peptide (S→E substitution) from tumour suppressor LKB1 was reported recently (PDB ID 4ZDR), the study resulted in a surprising and most likely unspecific binding mode for the peptide, different in two subunits of 14-3-3 [47, 48]. The reason for this may be the longer linker length or incomplete imitation of phosphorylation by a glutamate residue. The latter is also supported by the fact that, in the 4ZDR structure, the primary binding site is occupied by sulfate anion which apparently outcompeted the central Glu replacing the phosphoresidue. In this respect, the reported peptide conformation [47] should probably not be considered a genuine novel ligand-binding mode.

By contrast, in our study we validated conformations of phosphopeptides obtained in our chimeras by direct comparison of the crystal structures of the CH1 chimera with that of the 14-3-3σ complex with the synthesized HSPB6 phosphopeptide (PDB ID 5LU1). Importantly, with C_α_ r.m.s.d. of 0.23 Å, these completely different approaches provided almost identical structural information (Fig. 3C).

Interestingly, phosphopeptide binding within the 14-3-3 chimera CH1 resulted in protein compaction and significant increase in thermal stability, as evidenced by analytical SEC and fluorescence spectroscopy (Fig. 2), in line with partial stabilization of 14-3-3 by phosphate and phosphopeptides observed earlier [49]. The compaction and stabilization of chimeric 14-3-3 may be used to verify its correct assembly during production/purification prior to crystallization, since problems with phosphorylation and peptide binding could be immediately identified.

As an approach to facilitate structural studies on full 14-3-3 complexes, we propose that 14-3-3 chimeras with the optimized linker length (Fig. 1) can also be used for future construction of more complex, specific protein-protein 14-3-3 chimeras, if there is a single phosphorylated 14-3-3-binding site per chain, and if it is located upstream the rest of a 14-3-3 partner. For example, *ternary* complexes involving 14-3-3 scaffolds, which have long been discussed but are poorly evidenced so far, can now be more easily studied. To this end, fusion of two different phosphopartners to different 14-3-3 isoforms that are known to preferably heterodimerize [4-6] can pave the road for important structural studies. For some interactions, where binding of a protein/domain to 14-3-3 is possible only after a phosphopeptide is already bound in the AG, 14-3-3 chimeras can also be helpful. Examples include the ternary 14-3-3 complex, GF14c/OsfD1/Hd3a, regulating flowering in plants [50] or mammalian 14-3-3/HSPB6 regulatory complex, where binding of alpha-crystallin domain of HSPB6 most likely takes place after 14-3-3/phosphopeptide binding in the AG [27]. The modular principle of chimeras described in this study can be also generally adaptable for other phosphoserine/threonine binding proteins [51] and even beyond.

More practical applications of 14-3-3 chimeras include the perspective construction of various biosensors for *in vitro* and *in vivo* studies on their platform. Even before detailed structural characterization and optimization, successful Förster resonance energy transfer (FRET)-based biosensors for PKA [52] or MARK (microtubule affinity regulating kinase) [53], having the 14-3-3 core as part of the sensor, were reported. Conjugation of 14-3-3 chimeras with various reporters can benefit from the advent of various fluorescence lifetime imaging (FLIM)-based techniques which are significantly more precise and less background-dependent than FRET-based ones. Chemical modification of dedicated engineered Cys residue(s) with environmentally sensitive fluorescence labels may help to create valuable reporter systems for *in vitro* studies, chips and other diagnostic tools.

## EXPERIMENTAL PROCEDURES

### Cloning, expression and purification of 14-3-3 chimeras

Cloning, overexpression and purification of the monomeric mutant form of human 14-3-3ζ (14-3-3ζ_m_: ^12^LAE^14^ → ^12^QQR^14^) and the untagged C-terminally truncated human 14-3-3σ (14-3-3σΔC: residues 1-231) were described previously [27, 39, 40]. To facilitate crystallization of the protein, we previously followed the surface-entropy reduction (SER) approach [38] and cloned 14-3-3σΔC mutant Clu3 with ^75^EEK^77^ → ^75^AAA^77^ amino acid replacements into a modified pET28 vector containing a 3C-cleavable N-terminal hexahistidine tag [27]. cDNA of the 14-3-3 chimera with the HSPB6 peptide RRAS^16^APL (CH1) was obtained in one PCR step using the pET28-his-3C_14-3-3σΔC-Clu3 construct as a template by high-fidelity *Pfu* polymerase using T7-forward 5’-GACTCACTATAGGGAGACC-3’ and an excess of Clu3-B6p reverse primer 5’-ATATCTCGAG*TCA*CAACGGGGCGCTAGCGCGGCGCAGGGATCCCGATCCCGTCCACAGTGTCAG-3’ introducing the HSPB6 peptide and linker (GSGS) sequences and *XhoI* site. cDNA of the 14-3-3 chimeras with the Gli (CH2) or StARD1/BimEL (CH3) peptides were obtained on the basis of CH1 by the same procedure as for CH1 but using 5’-ATATCTCGAGTCATGCTTGAGCAGGATCACTAGCGCGGCGCAG-3’ or 5’-ATATCTCGAGTCAACGAGATCCCAGCAGGCTGCTGCGGCGCAGGGATC-3’ reverse primers, respectively, introducing the peptide and linker sequences and *XhoI* site. cDNA of CH1-CH3 was subsequently cloned into pET28-his-3C vector using *NdeI* and *XhoI* sites for restriction endonucleases and T4 DNA-ligase (SibEnzyme; www.sibenzyme.com). Correctness of all constructs was verified by DNA sequencing in Evrogen (www.evrogen.com).

All phosphorylated chimeras CH1-CH3 were obtained according to the identical scheme. Corresponding constructions in pET28-his-3C vector (kanamycin resistance) were used for co-transformation and co-expression in *E. coli* with a His-tagged catalytically active subunit of mouse PKA cloned in pACYC vector (chloramphenicol resistance) [27] under selection on both antibiotics. Protein overexpression in 1 L of Luria-Bertani media was induced at OD_600_ of 0.6 by addition of isopropyl-β-D-thiogalactoside to a final concentration of 0.5 mM for 20 h at 30 °C.

Purification was performed using subtractive immobilized metal-affinity chromatography (IMAC) and gel-filtration essentially as described [27]. Between IMAC1 and IMAC2 steps (loading/washing buffer (A): 20 mM Tris pH 8.0, 300 mM NaCl, 10 mM imidazole; elution buffer (B): buffer A with additional 500 mM imidazole) the chimeras were dialyzed to remove imidazole and simultaneously cleaved with 3C protease [27, 54] (1:1000 weight 3C:chimera ratio estimated by absorbance at 280 nm) resulting in target proteins with three extra residues GPH- at their N-terminus. The final polishing size-exclusion chromatography step was followed by immediate screening for crystallization conditions or biophysical characterization. The amount of protein obtained from 1 L of bacterial culture was usually enough to setup exhaustive initial screening and obtain diffraction quality crystals without further optimization. All final protein samples were homogenous according to a Coomassie-stained SDS-PAGE. Protein concentration was determined spectrophotometrically at 280 nm.

### Analytical size-exclusion chromatography

The oligomeric status and hydrodynamic properties of CH1 chimera, 14-3-3ζ_m_, and 14-3-3σΔC were assessed and compared using analytical SEC essentially as previously described [55]. 100 μL protein samples were pre-incubated for 30 min at room temperature and then loaded on a Superdex 200 Increase 10/300 column (GE Healthcare) equilibrated with a 20 mM Tris-HCl buffer, pH 7.6, containing 200 mM NaCl, 0.1 mM EDTA, and 3 mM β-mercaptoethanol (ME), at a flow rate of 1.5 mL/min, while monitoring absorbance at 280 nm. The column was calibrated by protein standards with known hydrodynamic radii that were used to determine average radii *R*_H_ of the species under study [55, 56]. Profiles were built using *Origin 9.0 Pro* software.

### Fluorescence spectroscopy

Intrinsic tryptophan fluorescence spectra of CH1 and 14-3-3σΔC (1.5 μM per monomer) were recorded using a Cary Eclipse fluorescence spectrophotometer (Varian Inc.) in a total volume of 600 μl on a 20 mM Hepes buffer, pH 7.1, 150 mM NaCl, 0.1 mM EDTA, 2 mM ME in the range from 305-450 nm upon excitation at 297 nm. The slits width was 5 nm and typically absorbance at the excitation wavelength was less than 0.1 to exclude the effects of inner filter. The spectra were registered in triplicate, averaged and then buffer-corrected.

To get insight into thermal stability of CH1 chimera in comparison with 14-3-3σΔC, we monitored changes in the intensity of fluorescence at 320 (*I*_320_) and 365 (*I*_365_) nm upon excitation at 297 nm as well as in light scattering at 90° angle (crossed monochromators 350/355 nm) during heating of the samples (0.7-7.7 μM concentration range) from 10 to 80 °C at a constant rate of 1 °C/min. Before the experiment the samples were equilibrated for 30 min at the initial temperature (10 °C). The ratio of *I*_320_(T)/*I*_365_(T) normalized from 0 to 100% represented the dependence of completeness of thermal transition, of unfolded fraction, on temperature and was used to estimate half-transition temperatures [42]. When possible, single wavelength was used to build analogous transition curves [56]. Graphs were built using *Origin 9.0 Pro* software.

### Crystallization and X-ray data collection

The 14-3-3 chimeras were subjected to crystallization screening immediately after purification by SEC using commercial screens including PACT, Procomplex (Qiagen), Index, Crystal Screen (Hampton Research) and JCSG+ (Molecular Dimensions). Sitting drops containing 200 nl protein at 10-23 mg/ml concentration (See Table 1) and 100-200 nl precipitant solution were set up in 96-well plates on a Mosquito robot (TTL Labtec). The crystals were difficult to optimize, however, in some cases random matrix microseeding appeared helpful (Table 1). The crystallizations were incubated at 20 °C and monitored using a plate Imager (Rigaku) equipped with Vis/UV-scanning and detection system.

X-ray diffraction data (Table 2) on small crystals grown directly in 96-well plates were collected at 100 K at I04 beamline of Diamond Synchrotron (UK) by using Pilatus detectors. Crystals were mounted on nylon loops and quickly cooled in liquid nitrogen, predominantly without using cryoprotection (See Table 1 for details). Two-three isomorphous crystals were obtained for each structure; the datasets with best resolution limits were used for structure solution and refinement.

### Crystal structure solution and refinement

Integration and scaling of diffraction data were done using *XDS/Xscale* [46] and *xdsme* [57]. Phasing of the CH1-CH3 was accomplished by molecular replacement in *MolRep* [58] using the dimer of the 14-3-3σ Clu3 mutant from the PDB ID 5LU1 as a search model. Initial attempts of phasing in the case of CH3 using a dimeric 14-3-3σ failed and could be possible only by using 14-3-3σ monomer as a search model, which gave three out of four subunits in the ASU, and the following manual building of α-helices in *Coot* [59] into the electron density map calculated with the phases from the initially found three 14-3-3 monomers, resulting in the fourth 14-3-3 subunit at a substantially more open protein conformation. The missing C-terminal fused phosphopeptides of chimeras and sulfate anions in the AG of 14-3-3 were manually added on the basis of difference electron density maps obtained after molecular replacement. Automated refinement in *Buster* 2.10.3 [60] initially included a rigid-body refinement of all chains and then an all-atom and individual B-factor refinement additionally restrained by the available non-crystallographic symmetry as well as by the ‘target’ high-resolution structures. Especially for CH3, as the ‘target’ structure we used a 14-3-3σ monomer (from PDB ID 5LU1). The quality statistics corresponding to the final models is presented in Table 2. The relatively higher R-factors in case of CH1X structure are more likely associated with the pronounced translational NCS detected in this crystal, which significantly complicated the refinement. In this case, *Zanuda* [61] was used to validate the *P* 2_1_ 2_1_ 2_1_ space group.

After the structures of CH1-CH3 had been established, we could use them to confirm that other isomorphous crystals had the same overall structure supporting the tracing and peptide conformation. All structural figures were prepared using Pymol 1.6.9 (Schrödinger). Atomic coordinates and structure factors have been deposited with the PDB under accession codes indicated in Table 2.

## ACKNOWLEDGMENTS

We are grateful to Vladimir Levdikov (YSBL, The University of York) for help with crystallization setups and to Sam Hart (YSBL, The University of York) for assistance with X-ray data collection. This research was supported by a grant from Russian Foundation for Basic Research 14-04-00146 to N.N.S., by the Program “Molecular and cell biology” of the Russian Academy of Sciences to N.N.S., and the Wellcome Trust grant 098230 to A.A.A..

Authors declare no conflict of interests.

